# Are Nonsense Alleles of *Drosophila melanogaster* Genes under Any Selection?

**DOI:** 10.1101/232090

**Authors:** Nadezhda A. Potapova, Maria A. Andrianova, Georgii A. Bazykin, Alexey S. Kondrashov

**Affiliations:** Institute of Information Transmission Problems (Kharkevich Institute) of the Russian Academy of Sciences, 127051, Moscow, Russia; Department of Bioengineering and Bioinformatics, Lomonosov Moscow State University, 119991, Moscow, Russia; Skolkovo Institute of Science and Technology, 143026, Moscow, Russia; University of Michigan, MI 48109, Ann Arbor, USA

**Keywords:** drosophila, loss-of-function, nonsense allele, negative selection, neutral evolution

## Abstract

A gene which carries a *bona fide* loss-of-function mutation effectively becomes a functionless pseudogene, free from selective constraint. However, there is a number of molecular mechanisms that may lead to at least a partial preservation of the function of genes carrying even drastic alleles. We performed a direct measurement of the strength of negative selection acting on nonsense alleles of protein-coding genes in the Zambian population of *Drosophila melanogaster*. Within those exons that carry nonsense mutations, negative selection, assayed by the ratio of missense over synonymous nucleotide diversity levels, appears to be absent, consistent with total loss of function. In other exons of nonsense alleles, negative selection was deeply relaxed but likely not completely absent, and the per site number of missense alleles declined significantly with the distance from the premature stop codon. This pattern may be due to alternative splicing which preserves function of some isoforms of nonsense alleles of genes.

## Introduction

A gene whose function does not contribute to fitness is on its way to becoming a pseudogene. Thus, with few exceptions (Xue et al. 2006; MacArthur et al. 2007), genes must be protected by negative, or purifying, selection which removes their loss-of-function (LoF) alleles. Nevertheless, genotypes of individuals carry substantial numbers of LoF alleles or, more precisely, of alleles that are likely to cause a complete loss of function of a protein-coding gene (MacArthur et al. 2012). Such alleles include nonsense substitutions which produce a premature stop codon as well as frameshift deletions, insertions, and complex mutations. In humans, there are 53-100 LoF alleles per genotype, including 21-27 nonsense alleles (MacArthur et al. 2012; Li et al. 2015). Data on nonsense alleles are more abundant and more reliable than on other LoF alleles, because calling frameshift alleles when genotypes are studied by resequencing is problematic. There are ~30 nonsense alleles per genotype in pig *Sus scrofa* (Groenen et al. 2012), ~100 in an alga *Chlamydomonas reinhardtii (Flowers et al. 2015)*, and ~18 in North American or ~35 in Zambian populations of *Drosophila melanogaster* (Lack et al. 2015; Yang et al. 2015).

Large per genotype numbers of nonsense and other LoF alleles may suggest that at least some of them do not, in fact, lead to a complete loss of function. Indeed, there is a number of molecular mechanisms that could ensure at least a partial preservation of function of an allegedly LoF allele, including alternative splicing, stop codon readthrough, and alternative translation initiation (Jagannathan and Bradley 2016). Several cases of functioning nonsense alleles have been described (Prieto-Godino et al. 2016).

Still, there is no doubt that, on average, a nonsense allele is more deleterious than a missense allele. For instance, a nonsense allele is three times more likely to lead to a disease than a missense allele (Krawczak et al. 1998). Per site prevalence of nonsense alleles in all studied populations is substantially lower than that of missense alleles (Mort et al. 2008; Yamaguchi-Kabata et al. 2008; Kono et al. 2016). Genes that harbor nonsense alleles have narrower expression profiles, are commonly involved in dispensable biological processes, and have many paralogs, which makes loss of their functions less deleterious (Lee and Reinhardt 2012; MacArthur et al. 2012; Yang et al. 2015).

To investigate the impact of nonsense alleles on the function of affected genes, we performed a direct measurement of the strength of negative selection acting within these alleles in a natural population of *D. melanogaster*.

## Materials and Methods

### D. melanogaster genome datasets

We used the DPGP3 dataset of genomes of Zambian *D. melanogaster* haploid embryos as our main dataset (Lack et al. 2015; http://www.johnpool.net/genomes.html). We used only those 196 genomes for which all the three major chromosomes, 2, 3 and X, were available. We also analyzed two smaller datasets (of ~50 individuals each) from Africa (AGES) and North America (NUZHDIN) (Lack et al. 2015). The annotation file has been downloaded from UCSC Genome Browser for version 3 of *D. melanogaster* genome (http://hgdownload.cse.ucsc.edu/goldenPath/dm3/database/flyBaseGene.txt.gz). Canonical splice variants of genes are from (http://hgdownload.cse.ucsc.edu/goldenPath/dm3/database/flyBaseCanonical.txt.gz). We used the longest isoform for every gene, in order to include as many nonsense mutations as possible in analysis.

### Data filtering

We focused on single-nucleotide nonsense substitutions. A gene was excluded from the analysis if the start codon was not ATG, if the stop codon was not TAA, TAG, or TGA, or if the length of the coding sequence was not in a multiple of 3. 90 genes that contain at least one nonsense allele with the frequency above 0.3 were excluded from estimates of negative selection, because such nonsense alleles are often spurious. Individual nonsense alleles located within the first or the last 5% of the length of the ORF were also excluded (MacArthur et al. 2012) from these estimates. 73% (1231/1689) of genes and 62% (1726/2786) of nonsense alleles survived this filtering.

### pN/pS estimation

pS was calculated using 4-fold degenerate synonymous sites, and pN was calculated from non-degenerate sites at second positions within each codon only. For each site of the corresponding category, site-specific pN or pS were calculated as

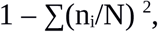

where N is the number of genotypes, n_i_ is the number of genotypes carrying a certain nucleotide, and summation is over all four nucleotides. pN and pS were then obtained by averaging these values over all sites of the corresponding category.

To calculate pN and pS for nonsense alleles, we used only non-singleton nonsense alleles (i.e., those observed two or more times in our sample). We analyzed only those synonymous and missense polymorphisms that were nested within the nonsense alleles (Figure 1), because when a polymorphism is present in all nonsense alleles there is a chance that it originated before the nonsense mutation.

**Fig. 1.**
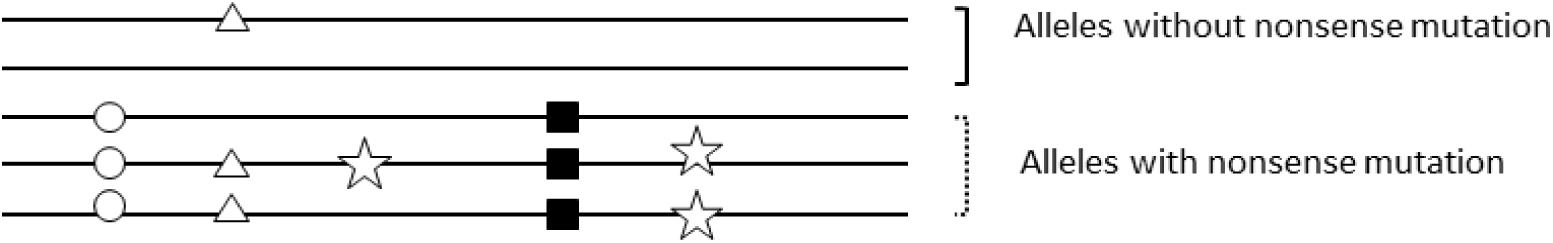
Schematic representation of mutation types in nonsense alleles. The presence of a nonsense mutation (shown as a square) subdivides the sample into nonsense and sense alleles. For analysis of pN/pS, we considered only those synonymous or missense mutations that only occurred in nonsense alleles, but did not occur in all nonsense alleles (stars). Such mutations are most likely to have arisen after the nonsense mutation against its background. Mutations in all nonsense alleles (circles), or mutations occurring in some sense alleles (triangles), were not considered.

Calculation of pN/pS ratios for mutations nested within frequency-matched synonymous alleles with those nested within nonsense alleles was performed in genes with nonsense alleles using the formula described above.

The confidence intervals for pN/pS ratios were estimated from 10,000 bootstrap trials resampling case. Bootstrapping was performed by individual genes.

pN and pS estimations for Figure 5 were calculated for sliding windows of width 210 (70 codons), with the step 50.

### *D. melanogaster* RNA-seq datasets

We used RNA-seq dataset SRR3135045 (von Heckel et al. 2016) for Zambian *D. melanogaster* from the NCBI SRA (https://www.ncbi.nlm.nih.gov/sra). Raw reads were downloaded with SRA Toolkit (v. 2.8.0). Then we trimmed this data using Trimmomatic (v. 0.32) and made quality control using FastQC (v. 0.10.1). Transcriptome was mapped to *D. melanogaster* reference genome (dm3) with TopHat (v. 2.1.0). Coverage for nonsense exons was calculated using BEDTools (v. 2.16.2) with option “coverage -counts -abam”.

Then for each gene, we calculated the relative density of nonsense exon reads as the following ratio:

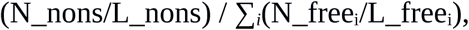

where N_nons is the number of reads mapped onto an exon carrying the nonsense mutation, L_nons is the length of this exon, N_free_i_ is the number of reads mapped onto the ith nonsense-free exon, L_free_i_ is the length of the *i*th exon. The final value was obtained by averaging over all genes.

## Results

### Prevalence of nonsense alleles

We investigated only nonsense alleles that resulted from a single nucleotide substitution. Below, the term ‘nonsense allele’ refers to both a nonsense mutation and a haplotype which carries it.

In 196 haploid genotypes of Zambian *D. melanogaster* we detected, within canonical isoforms of 13,300 protein-coding genes, 1,726 nonsense alleles within 1,231 genes. Among these genes, 767 carried only singleton nonsense alleles, i.e. nonsense alleles that were observed in just a single genotype. The remaining 464 genes carried both singleton and non-singleton or only non-singleton nonsense alleles. The total number of singleton nonsense alleles was 1,236. On average, each genotype contained 35 genes with 36 nonsense alleles (including singletons), or 30 genes with 31 nonsense alleles (excluded singletons). The proportions of genes that harbor nonsense alleles were similar between chromosomes 2 and 3, but twice as low for X chromosome (Supplementary Table 1), in line with higher efficiency of negative selection against nonsense alleles in hemizygous state (Mackay et al. 2012).

We observed an excess of nonsense alleles near the 5′- and, especially, the 3′-end of genes (Figure 2), where they may not always destroy the function completely (Wetterbom et al. 2009; Lee and Reinhardt 2012).

**Fig. 2.**
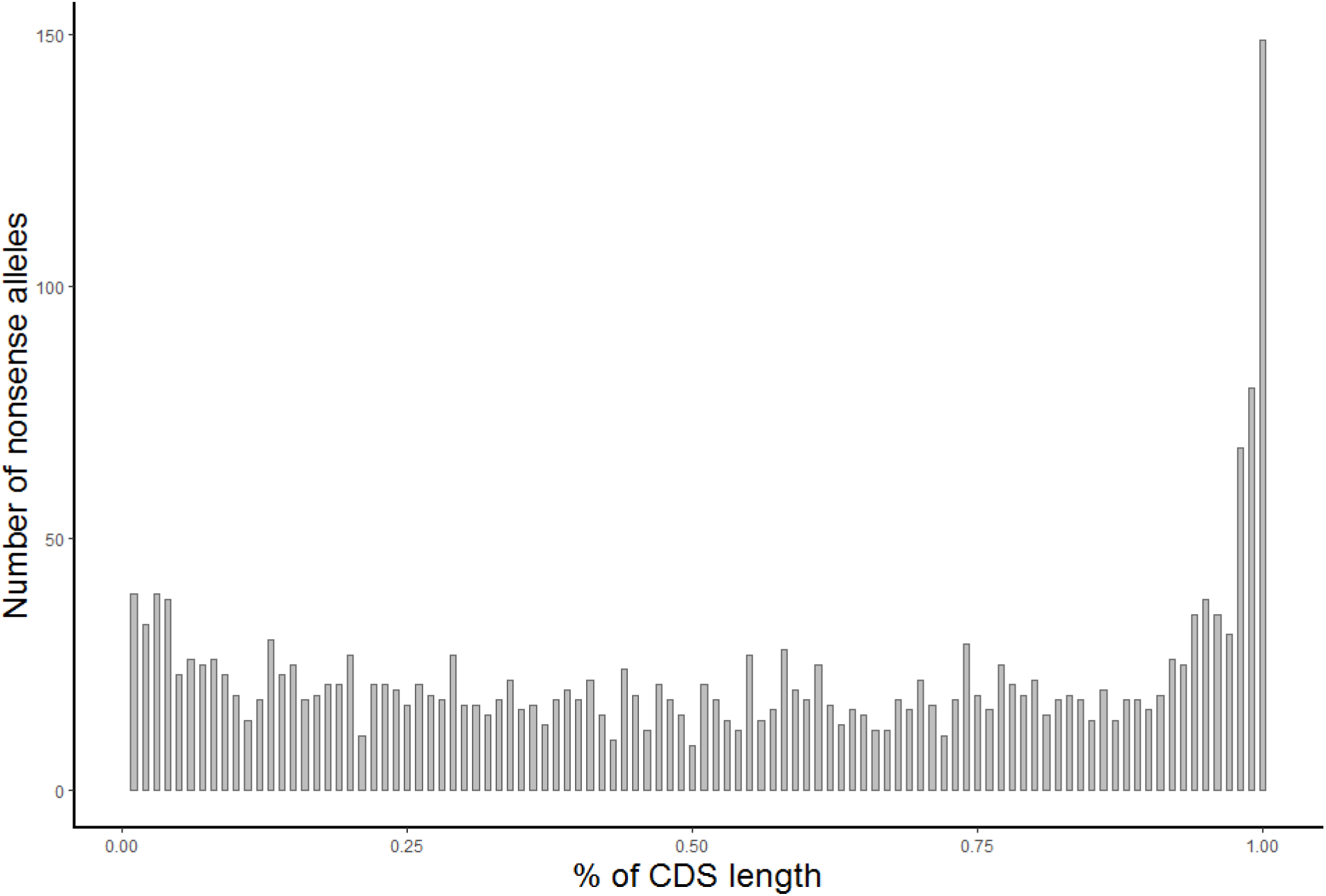
Relative positions of nonsense mutations within coding portions of genes which harbor them.

Figure 3 presents the distribution of frequency x of nonsense alleles. In agreement with the data obtained previously (Li and Stephan, 2006), we see a substantial excess of very rare nonsense alleles (i. e., of singletons and of those that appeared twice in our sample of 196 genotypes) over the neutral expectation of ~1/x (Wright 1931; Kimura 1983), which must be at least partially due to negative selection against them.

**Fig. 3.**
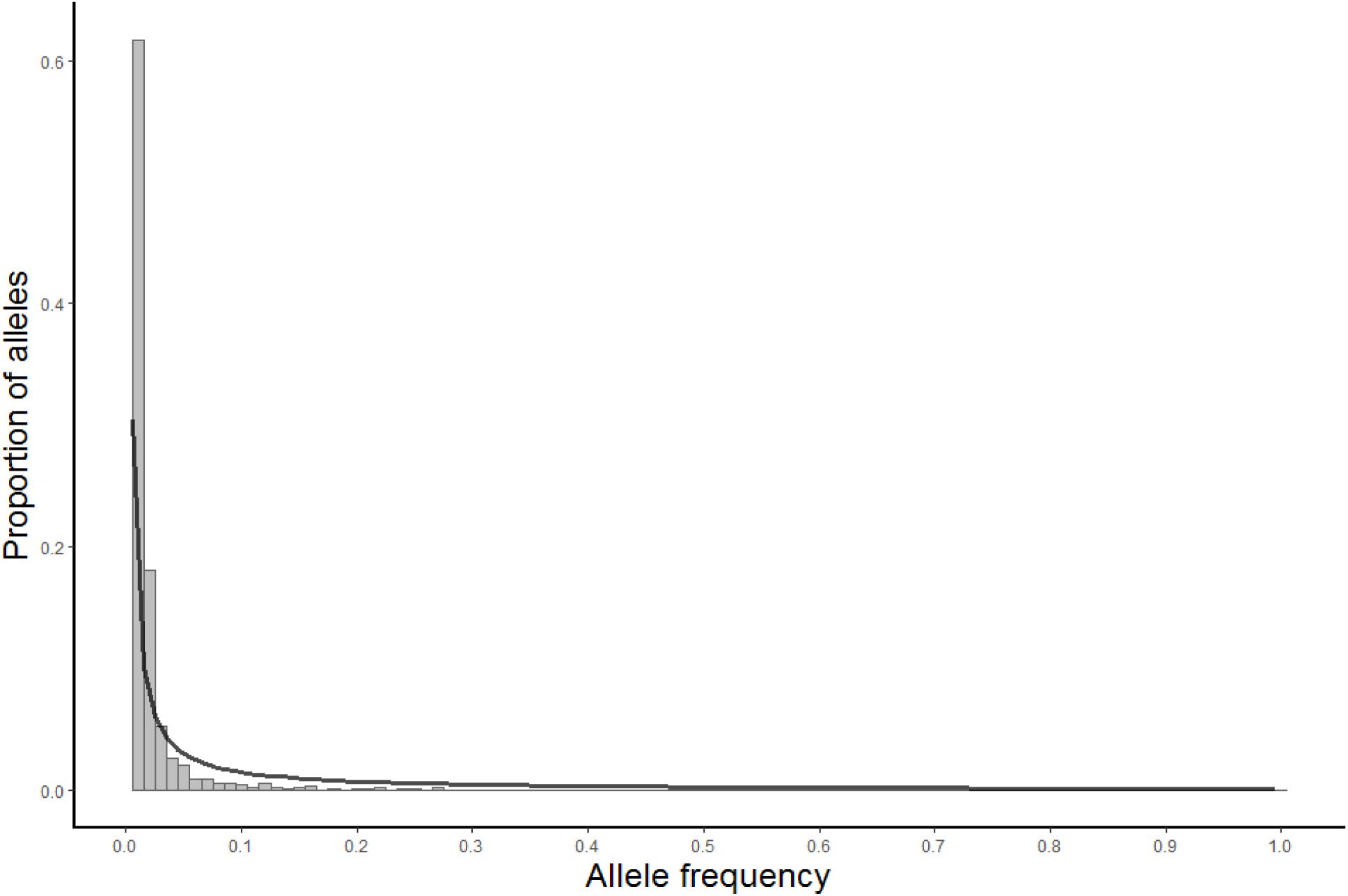
The observed distribution of frequencies of nonsense (grey bars) alleles and the ~1/x expected distribution (black line).

### Negative selection in nonsense alleles

Next, we asked whether selection affects polymorphisms that segregate at the genetic background of a nonsense allele. First, we compared the numbers of missense and synonymous SNPs in genes with and without nonsense alleles. After that, we compared these numbers in nonsense-carrying vs. nonsense-free alleles of genes that possess non-singleton nonsense alleles (Figure 4).

**Fig. 4.**
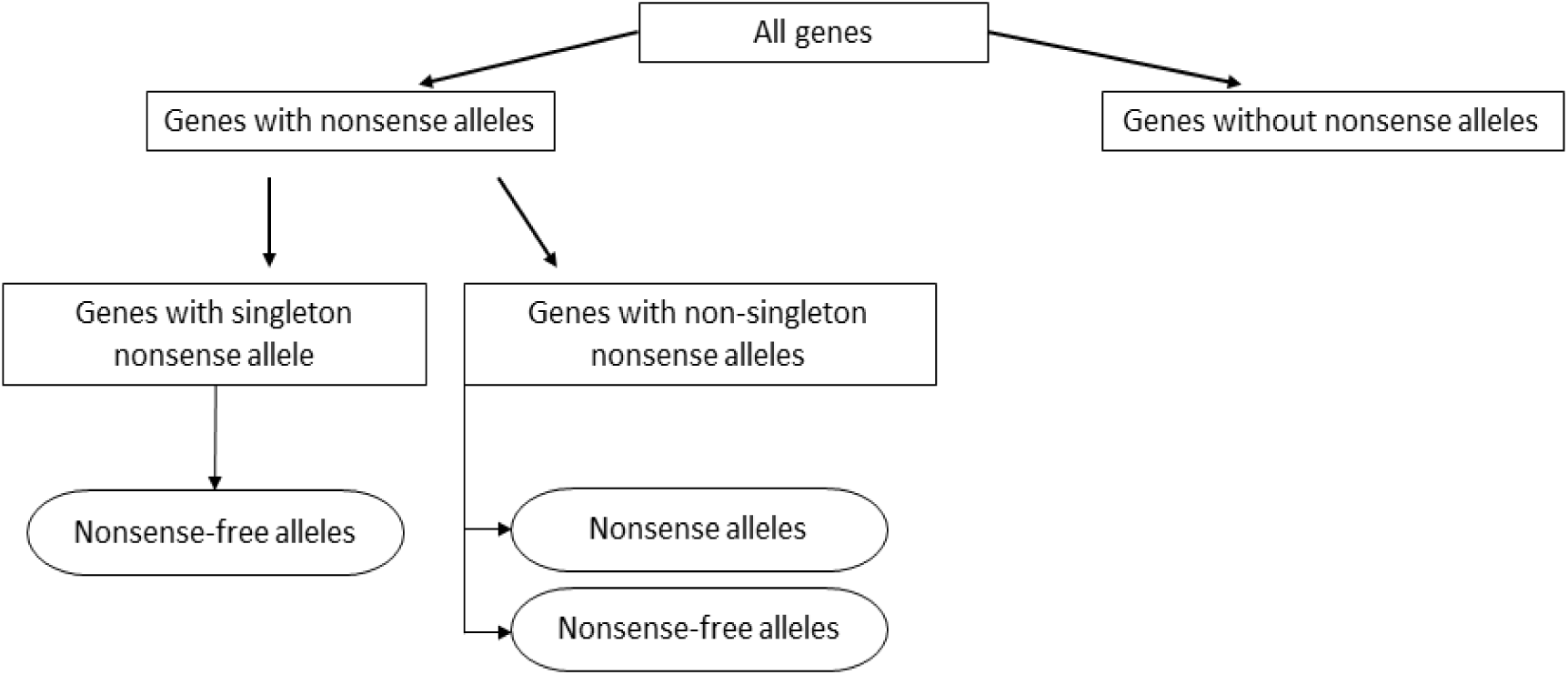
Workflow of data analysis. Classes of genes are shown in rectangles and classes of alleles are shown in ovals.

Table 1 presents data on the strengths of negative selection, characterized by pN/pS ratios, in classes of genes defined by the presence, within our sample of genotypes, of nonsense alleles in them. Not surprisingly, genes that carry nonsense alleles, and especially non-singleton nonsense alleles, are, on average, under weaker selection. Values for pN/pS calculated separately only for those alleles of genes that do not carry nonsense mutations are only slightly lower than the values for all alleles of the same genes, because nonsense alleles are rare. By contrast, the value of pN/pS obtained for non-singleton nonsense alleles on the basis of missense and synonymous polymorphisms nested within them is much higher than that for alleles that do not carry a nonsense mutation, and is not significantly different from 1, indicating that selection against missense mutations is reduced or absent. Such polymorphisms are present in only 140 out of the 464 genes that harbor non-singleton nonsense alleles, leading to wide confidence interval for this value (pN and pS values separately are shown in Supplementary Tables 2, 3). Still, the same analysis using smaller data sets of other *D. melanogaster* populations shows similar patterns (Supplementary Tables 4, 5).

**Table 1.** Average pN/pS in Zambian population of *D. melanogaster\**.

|  | All 12842 genes | 11611 genes without nonsense alleles | 767 genes with only singleton nonsense alleles | 464 genes with only non-singleton nonsense alleles |
| --- | --- | --- | --- | --- |
| All alleles | 0.106<br>[0.103-0.109] | 0.093<br>[0.091-0.095] | 0.186<br>[0.171-0.203] | 0.262<br>[0.238-0.290] |
| Non-nonsense alleles | 0.106<br>[0.103-0.109] | 0.093<br>[0.091-0.095] | 0.186<br>[0.171-0.202] | 0.256<br>[0.231-0.283] |
| Polymorphisms nested within 567 non-singleton nonsense alleles | - | - | - | 0.803<br>[0.584-1.121] |

\*95% CIs are shown in square brackets.

We analyzed Zambian *D. melanogaster* transcriptome data and found that the ratio of the densities of reads for nonsense-carrying over nonsense-free exons is 0.39 [95% CI: 0.32 – 0.46], indicating that a substantial proportion of nonsense-carrying exons are incorporated only in rare splice isoforms. Table 2 presents data on pN/pS ratios for polymorphisms nested within non-singleton nonsense alleles separately for one-exon genes, for exons of multi-exon genes that do not carry a nonsense mutation, and for exons of multi-exon genes that carry a nonsense mutation.

**Table 2.** Average pN/pS for polymorphisms nested within nonsense alleles*.

| pN/pS | 121 nonsense alleles<br>of 99 one-exon genes | 446 nonsense alleles of 335 multi-exon genes |  |
| --- | --- | --- | --- |
|  |  | Exon with nonsense<br>mutation | Exons without<br>nonsense mutation |
|  | 1.020 [0.770-1.530] | 0.906 [0.556-1.500] | 0.787 [0.459-1.359] |

\*95% CIs are shown in square brackets.

Figure 5 shows how the values of pN and pS, calculated for nested polymorphisms only, depend on the distance from the premature stop codon along the coding sequence of the gene. pN decreases with distance from the stop codon (the slope of the linear trend: −2.3 × 10^−7^ [95% CI: −3.9×10^−7^ – −5.9x10^−8^], p=0.01787), while pS does not change (the slope is 3.9×10^−8^, with 95% CI: −3.8×10^−7^ – 4.6×10^−7^, p=0.8186).

**Fig. 5.**
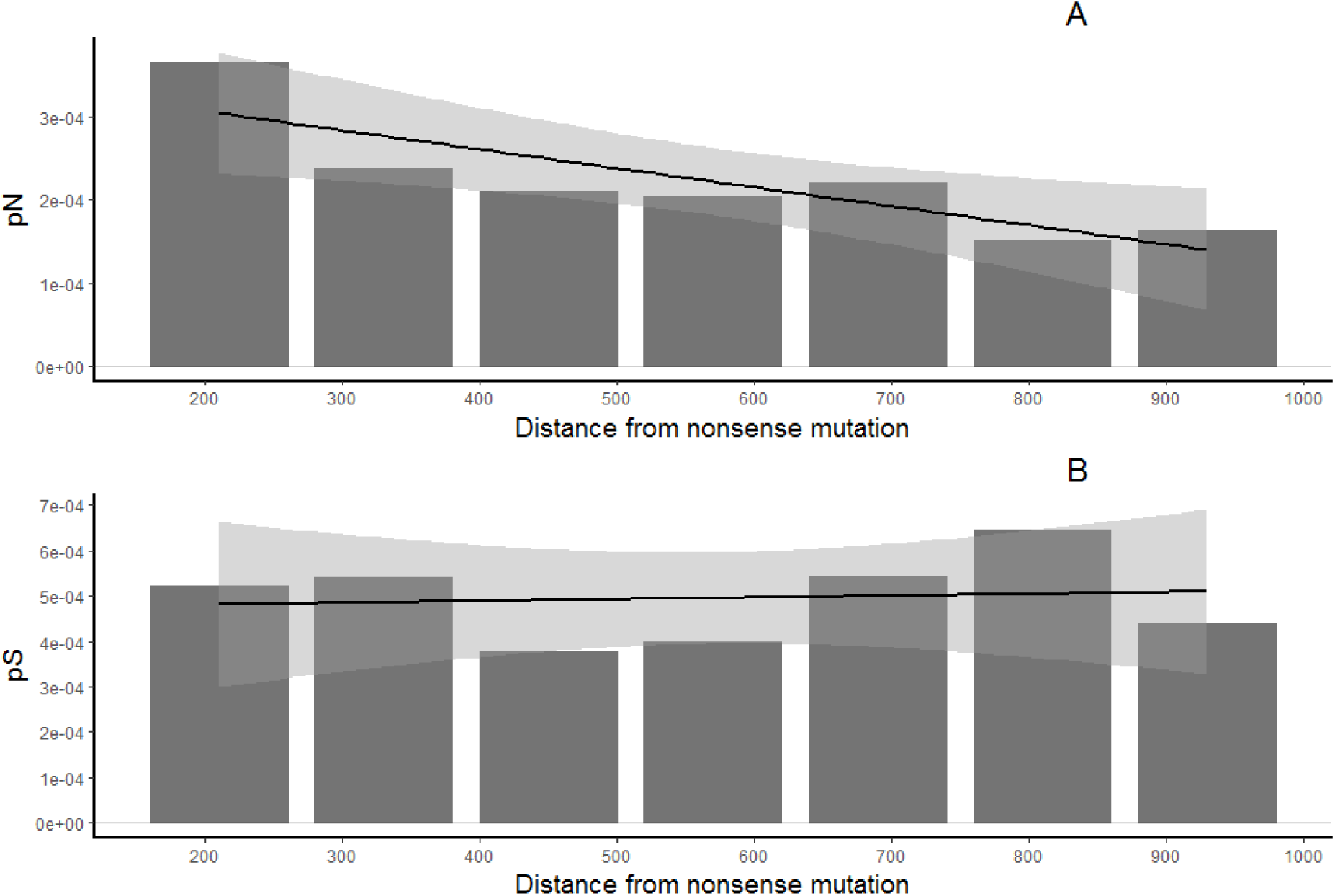
Dependencies of pN (A) and pS (B) in nonsense alleles on the distance along the CDS (introns excluded) from the nonsense mutation. Black lines show the linear trend with 95% CI indicated by shading.

## Discussion

Our main goal was to determine whether nonsense mutation-carrying alleles of genes retain some residual function and, thus, remain under some negative selection. We have found that within the exon of a nonsense allele which carries a premature stop codon, the pN/pS ratio, calculated on the basis of missense and synonymous mutations that are nested within the nonsense allele and therefore likely appeared after the nonsense mutation, is not significantly different from 1, indicating total relaxation of selection (Table 2). For other exons of nonsense alleles, we obtained a slightly lower value of pN/pS, which, however, is not significantly different from the first one or from 1.

pN, but not pS, declines with the distance from a premature stop codon (Figure 5). This contrast suggests that selection plays a role in the decline of pN. There could be two not mutually exclusive causes for this pattern. First, residual negative selection probably operates on exons that do not carry a premature stop codon, because some of them are not incorporated into all isoforms produced by alternative splicing. Second, even if negative selection is absent throughout the whole nonsense mutation-carrying allele, the observed effect could be due to its recombination with functional alleles depleted of missense substitutions. Unfortunately, our data are insufficient to discriminate between these two possibilities.

Nonsense alleles are mostly rare, so that missense and synonymous mutations nested within them are even rarer. In fact, all such mutations that are present in the data we used are singletons. Because negative selection leads to an excess of rare alleles, pN/pS ratios calculated on the basis of such mutations must be inflated, even without any relaxation of selection. We investigated this by calculating the pN/pS ratios for mutations nested within frequency-matched synonymous alleles with those nested within nonsense alleles. As expected, the average of these ratios (0.611 [0.556-0.770]) is much higher than for all mutations; however, it was still significantly below 1. Thus, relaxation of negative selection acting on nonsense alleles of genes appears to be real.

Analyses of small samples of genotypes from two other populations produced results similar to those reported above, but with even wider confidence intervals (Supplementary Table 4, 5). Obviously, it would be very desirable to analyze a much larger sample of genotypes. A dataset of more than 1,000 Drosophila genotypes is available (Lack et al. 2015), but they are of multiple geographical origins, so that using the values of pN/pS from this dataset as proxies for negative selection is problematic. Unfortunately, the already available massive data on human diploid genotypes are not easy to use for our purposes, because it is not always possible to distinguish maternal and paternal sequences. Hopefully, larger sets of haploid genotypes from the same population, which in the case of *Drosophila* can be obtained either from haploid embryos or from inbred lines, will soon become available.

Overall, we investigated the possibility of some residual negative selection acting on nonsense alleles of protein-coding genes of *D. melanogaster*. Our results are consistent with complete relaxation of selection within those exons that carry premature stop codons. However, there may be some weak negative selection within other exons, possible due to alternative splicing of the nonsense-containing exon.

## Acknowledgements

We thank Anna Klepikova and Michail Schelkunov and all laboratory team for helpful comments and useful discussions for this article. This work was supported by the Russian Science Foundation (grant number 14-50-00150).

